# Biguanide-PROTACs: Modulating Mitochondrial Proteins in Pancreatic Cancer Cells

**DOI:** 10.1101/2024.03.17.585436

**Authors:** Julie Vatté, Véronique Bourdeau, Gerardo Ferbeyre, Andreea R. Schmitzer

**Author notes:** Département de chimie, Faculté des arts et des sciences, Université de Montréal. 1375 a. Thérèse Lavoie-Roux, Montréal, Québec, Canada H2V 0B3. (ARS).

## Abstract

This study focuses on the synthesis of Biguanide-PROTACs, formed by conjugating the biguanide motif with diverse E3 enzyme ligands and spacers. Evaluation of their activity on pancreatic cancer cell (KP4) proliferation established a correlation between membrane permeability and median effective concentration. Mechanistic insights revealed that only two compounds exhibited biguanide-like AMPK activation, while only one hydrophobic compound uniquely altered mitochondrial protein levels. The prospect of developing and expanding the Biguanide-PROTAC library holds promises, offering potential insights into biguanide mechanisms and the creation of more potent anticancer agents. This study contributes to understanding the intricate interplay between compound structure, permeability, and anticancer activity, paving the way for targeted drug development in pancreatic cancer treatment.

## Introduction

Numerous studies have investigated the impact of biguanidium salts, often exemplified by metformin and phenformin (Fig 1A), on the proliferation of various cancer cell types, with a particular focus on their potential in treating pancreatic cancer.(1-8) The challenges posed by late-stage detection have rendered this cancer difficult to manage due to its advanced progression.(9) Furthermore, surgical or radiation interventions are constrained by the intricate anatomical proximity of the tumour to surrounding organs, limiting treatment options primarily to chemotherapy. Recognizing the imperative for novel therapeutic approaches targeting pancreatic cancer, derivatives of biguanides have emerged as promising candidates. These compounds have demonstrated efficacy across diverse pancreatic cell lines and in animal models, exhibiting notable absence of toxicity or adverse effects.(10-15) Nevertheless, a comprehensive understanding of their mechanism of action is an ongoing area of intensive investigation. Unravelling the correlation between their anticancer properties and chemical structure is crucial for refining the design of more potent agents to enhance their therapeutic potential.(16-19) The antiproliferative activity of biguanidium salts has been attributed to their ability to selectively target mitochondria, owing to their cationic nature. This targeting mechanism culminates in the inhibition of oxidative phosphorylation (OXPHOS) and activation of the AMPK pathway, a key cellular signalling pathway.(20-28) Metformin, a prominent member of this class, is implicated in modulating Complex I of the respiratory chain, inhibiting the oxidation of NADH into NAD^+^ and potentially reducing ATP production as a consequence.(29-31) Notably, this effect may also involve interactions with other constituents of the electron transport chain (ETC), such as the glycerol-3-phosphate shuttle.(32, 33) Despite these insights, certain studies posit that biguanides exhibit no discernible activity on isolated mitochondria.(34) This intriguing observation underscores that their mode of action is contingent upon the cellular localization, suggesting that an accumulation of the drug within mitochondria is requisite for inducing the inhibition of oxidative phosphorylation.(35) This could imply a multifaceted interaction between biguanides and mitochondrial components, warranting further exploration to unravel the intricacies of their cellular effects.

**Fig 1.**
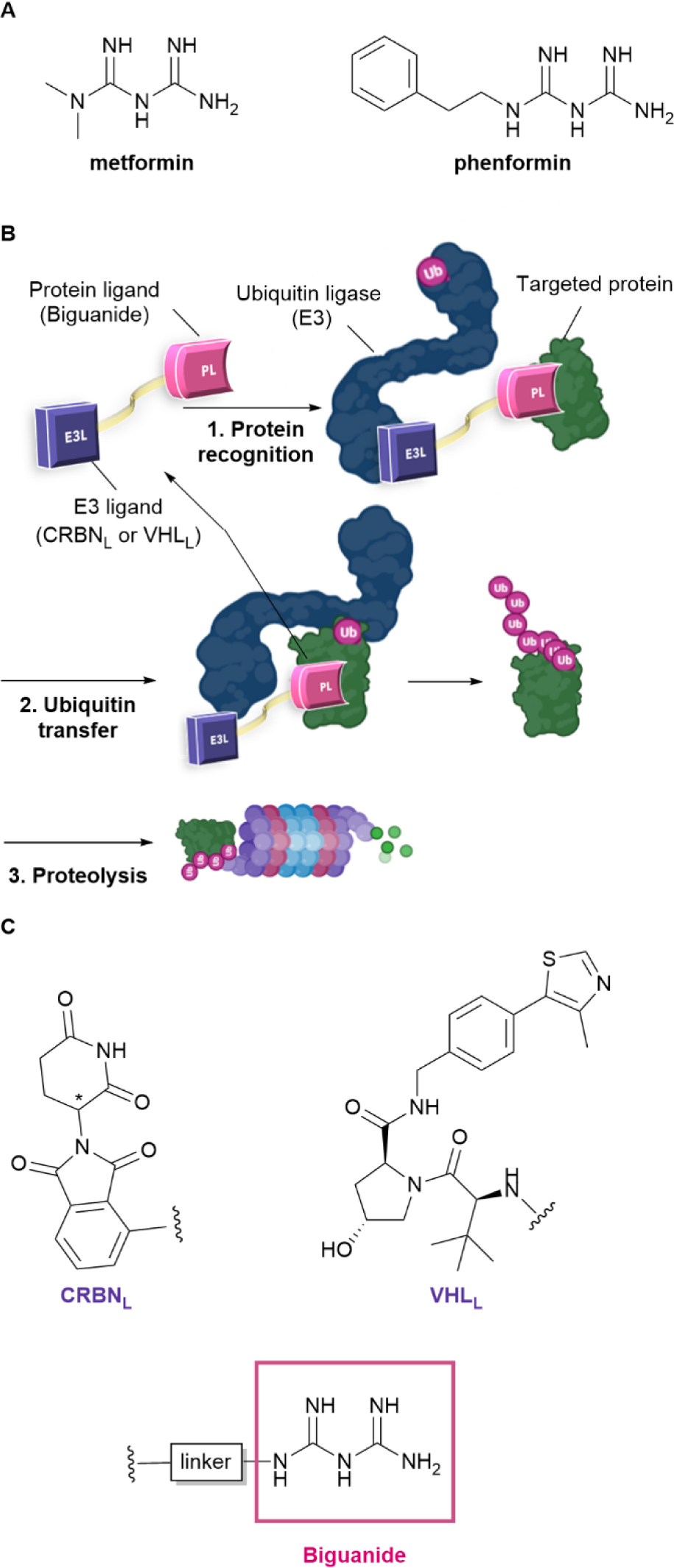
Design and mechanism of Biguanide-PROTAC. (A) Structures of metformin and phenformin. (B) Mode of action of the PROTAC compound leading to proteolysis. (C) Design and structure of Biguanide-PROTACs derivatives targeting the enzyme CRBN or VHL.

To delve deeper into the mechanisms of biguanides and pave the way for the development of novel anticancer agents, we have synthesized Biguanide-PROTACs (Proteolysis Targeting Chimeras) (Fig 1B).(36) These innovative molecules possess a bifunctional structure, comprising two distinct pharmacophores linked by a spacer. The first pharmacophore is designed to target a specific protein of interest, while the second binds to an ubiquitin ligase E3, thereby triggering the degradation of the targeted protein.(37) This unique compound is expected to form a ternary complex, bringing its two binding proteins in close proximity. Subsequently, the transfer of ubiquitin from the E3 enzyme to the protein of interest facilitates its recognition and degradation by the proteasome. This process liberates the PROTAC molecule, allowing it to bind to another protein and initiate a new cycle of degradation.(38) Unlike conventional inhibitors, which require a stoichiometric quantity of ligand to inhibit a protein’s action, PROTACs can be introduced in catalytic quantities. A single molecule of PROTAC has the potential to induce the degradation of multiple proteins. The development of such molecules holds the promise of enhancing the efficacy of biguanide derivatives by facilitating the targeted degradation of their specific protein targets. This approach represents a significant departure from traditional inhibition strategies. Moreover, it’s noteworthy that ubiquitin ligase inhibitors, also referred to as E3 ligands, have established themselves as effective anticancer agents in their own right.(39) This underscores the potential synergy between the targeted protein degradation facilitated by Biguanide-PROTACs and the established efficacy of E3 ligand-based anticancer therapies, presenting a multifaceted avenue for advancing cancer treatment strategies.

In addition to potentials in cancer treatments, PROTACs serves as valuable tools for investigating the interaction between a pharmacophore and a targeted protein, as highlighted in the literature.(40) In this context, the development of a Biguanide-PROTACs presents an opportunity to experimentally validate the reported interaction between the biguanide moiety and proteins from the respiratory chain of mitochondria. This validation is achieved through the observation of the targeted protein degradation. In pursuit of this goal, we meticulously designed six Biguanide-PROTACs, connecting a biguanide moiety to two distinct E3 ligands via various linkers. These ligands were aptly named CRBN_L_ (ligand for the Cereblon enzyme) and VHL_L_ (ligand for the Von Hippel Lindau protein) (Fig 1C). The design of these bifunctional molecules was guided by considerations of the linker’s nature and size, as these factors play a pivotal role in facilitating ternary complex formation and determining the efficacy of PROTACs.(41)

## Results and Discussion

### Design and Synthesis of Biguanide-PROTAC Derivatives

A range of primary amines underwent condensation with dicyandiamide in the presence of trimethylsilyl chloride, resulting in the synthesis of corresponding biguanide derivatives with yields ranging from 70% to 89% (see Supporting Information). Compounds **1, 5, 6** and **7** were subsequently engaged in diverse reactions, such as peptide coupling and copper-catalyzed coupling to incorporate the biguanide motif into a library of final compounds. Incorporation of a phenylethynylbenzyle biguanide motif, previously investigated within our research group, further enriched the compound library.(15) This particular motif, known for its intriguing biological properties, especially its ability to traverse phospholipid membranes, served as a pivotal building block (compound **5**) for the synthesis of the desired PROTACs. The synthetic routes for developing Biguanide-PROTACs recruiting CRBN (**11, 12, 16**, and **20**) and VHL (**29** and **32**) are depicted in Scheme 1, with detailed information on the preparation of reactants available in the Supporting Information. The exploration of diverse synthetic pathways involves the independent synthesis of E3 ligands and biguanide derivatives. Each fragment, distinguished by functionalities such as amine, carboxylic acid, azide, or alkyne, exhibits specific roles, allowing them to react synergistically, ultimately yielding the final compounds. Crucially, the linker within these structures can be subject to modification by introducing additional units with complementary functions. This strategic design enables the easy extension of the Biguanide-PROTACs library, fostering versatility and adaptability and also facilitating the fine-tuning of these molecules to optimize their pharmacological properties.

**Scheme 1.**
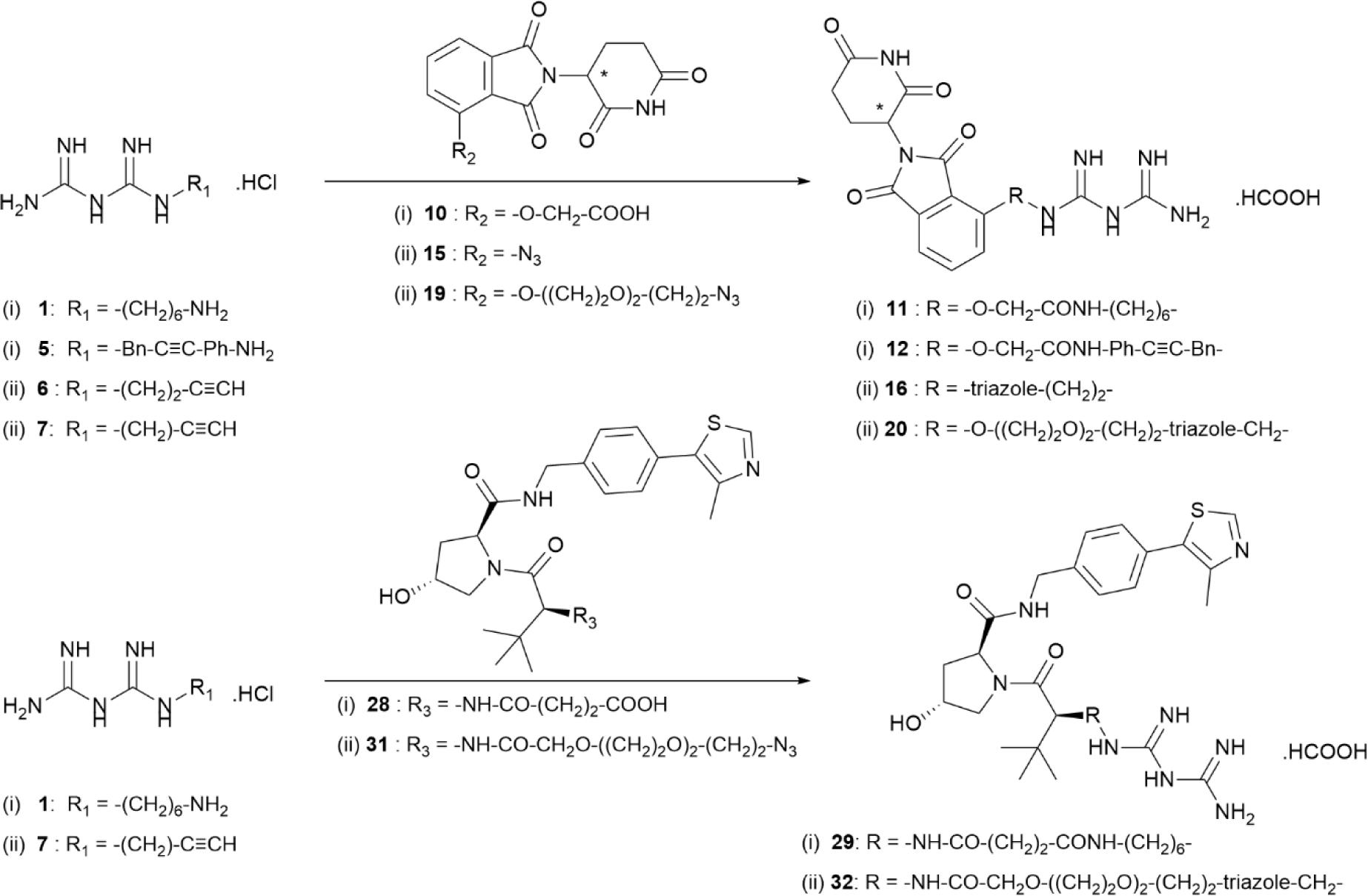
Synthesis of PROTAC-Biguanide compounds based on the CRBN or the VHL ligand. (i) HATU, DIPEA, DMF, r.t, 5-12 h, (**11**: 34%; **12**: 6%; **29**: 7%); (ii) CuI, Na-L-Ascorbate, DMF, r.t, 3-16 h (**16**: 13%; **20**: 14%; 32 : 29%).

### *In vitro* Antiproliferative Activity

In order to evaluate the antiproliferative efficacy of the synthesized Biguanide-PROTACs, a 72-hour incubation with various concentrations of these molecules was conducted using a pancreatic cancer cell line (KP4). The concentration required to reach the half maximal concentration affecting growth and viability (EC_50_) was determined for each compound and compared with that of metformin, a widely recognized antiproliferative agent. The antiproliferative properties of all tested compounds were evident, as manifested by their characteristic sigmoidal growth inhibition curves. Interestingly, while compounds **16** and **20** exhibited comparable efficacy to metformin, the majority of the designed PROTACs demonstrated a superior performance, necessitating a lower concentration than metformin to achieve a significant reduction in cell growth and viability (Fig 2).

**Fig 2.**
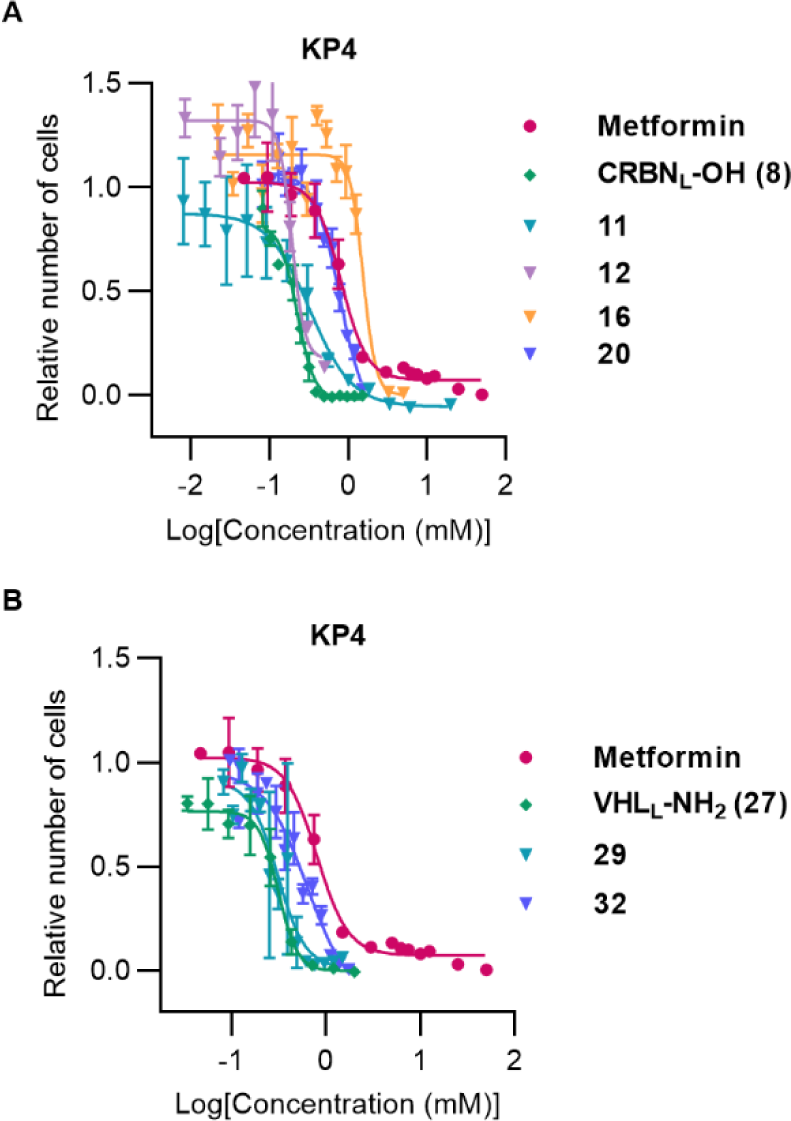
Dose-response curves of compounds Metformin, CRBN and VHL ligand derivatives in KP4 cells. Relative number of cells (%) was determined with crystal violet cytotoxicity assay over 72 h.

Prior to conjugation with the biguanide moiety, the CRBN and VHL ligands (designated as **8** and **27**, respectively; see Supporting Information) underwent testing to assess their individual antiproliferative activities. These ligands, represented by the green curves (Fig 2), exhibited superior antiproliferative effects compared to the CRBN_L_-Biguanide and VHL_L_-Biguanide conjugates. This intriguing observation suggests that the hybridization of E3 ligands with biguanides might influence their binding with the respective enzymes, impacting pharmacodynamics. Additionally, the alteration in physicochemical properties resulting from this conjugation could play a role in affecting pharmacokinetics. The net effect is an intermediate activity observed in the PROTACs, positioning them between the antiproliferative activities of the two parent pharmacophores. The observed relative activity was systematically correlated with the chemical structure of each compound to elucidate a structure-activity relationship. In this endeavour, partition coefficients (cLogP) were calculated to compare the hydrophobicity of the synthesized PROTACs and their respective controls, aiming to establish a discernible trend between their permeability in biological membranes and EC_50_ values (Table 1).

**Table 1.**
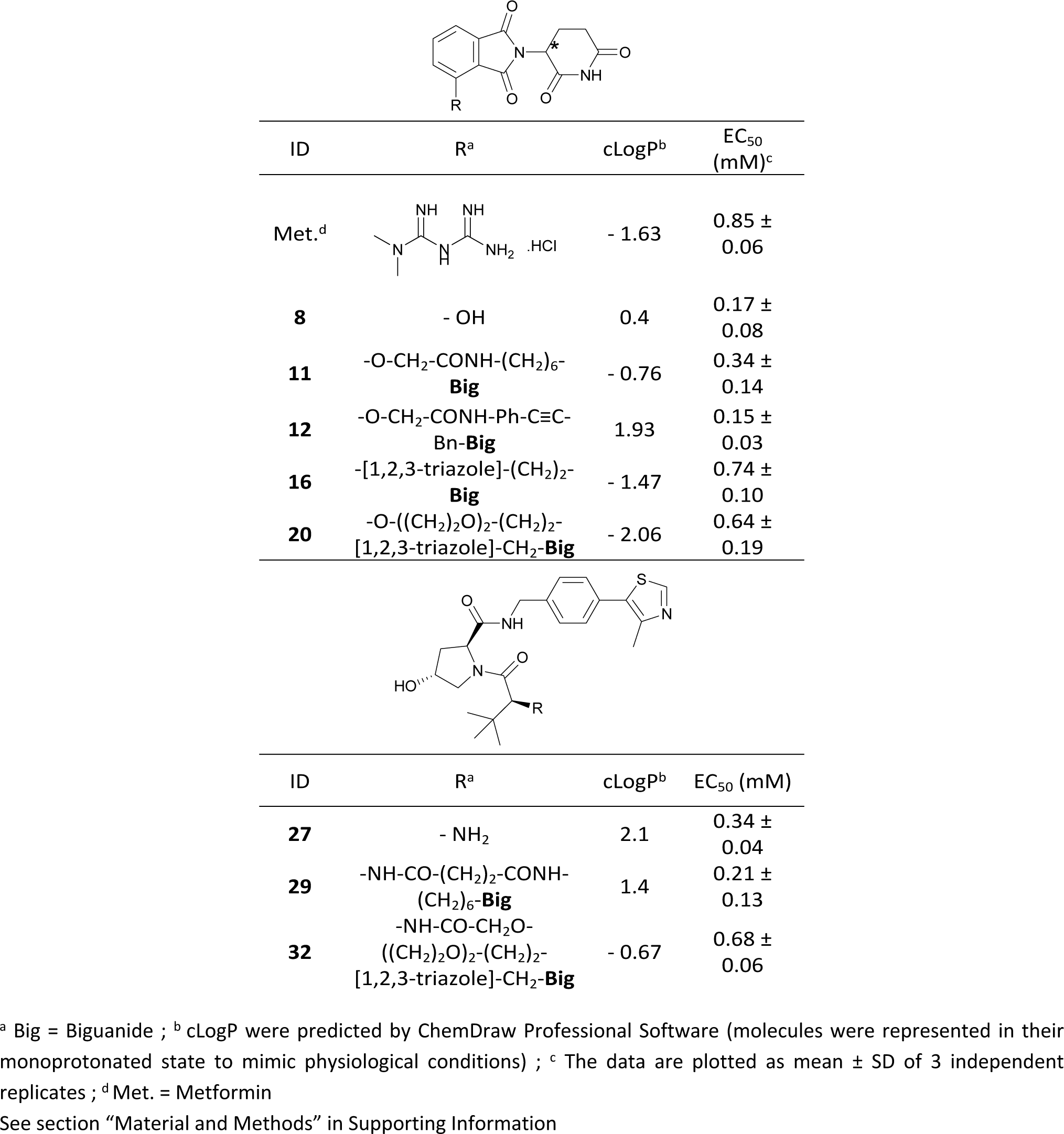
EC_50_ of Biguanide-PROTAC derivatives in KP4 cells after 72 h of treatment (mean ± SD)

|  |  |  |  |
| --- | --- | --- | --- |
| ID | R<sup>a</sup> | cLogP<sup>b</sup> | EC<sub>50</sub> (mM)<sup>c</sup> |
| Met.<sup>d</sup> | .HCl | - 1.63 | 0.85 ± 0.06 |
| **8** | - OH | 0.4 | 0.17 ± 0.08 |
| **11** | -O-CH<sub>2</sub>-CONH-(CH<sub>2</sub>)<sub>6</sub>-**Big** | - 0.76 | 0.34 ± 0.14 |
| **12** | -O-CH<sub>2</sub>-CONH-Ph-C≡C-Bn-**Big** | 1.93 | 0.15 ± 0.03 |
| **16** | -[1,2,3-triazole]-(CH<sub>2</sub>)<sub>2</sub>-**Big** | - 1.47 | 0.74 ± 0.10 |
| **20** | -O-((CH<sub>2</sub>)<sub>2</sub>O)<sub>2</sub>-(CH<sub>2</sub>)<sub>2</sub>-[1,2,3-triazole]-CH<sub>2</sub>-**Big** | - 2.06 | 0.64 ± 0.19 |
| ID | R<sup>a</sup> | cLogP<sup>b</sup> | EC<sub>50</sub> (mM) |
| **27** | - NH<sub>2</sub> | 2.1 | 0.34 ± 0.04 |
| **29** | -NH-CO-(CH<sub>2</sub>)<sub>2</sub>-CONH-(CH<sub>2</sub>)<sub>6</sub>-**Big** | 1.4 | 0.21 ± 0.13 |
| **32** | -NH-CO-CH<sub>2</sub>O-((CH<sub>2</sub>)<sub>2</sub>O)<sub>2</sub>-(CH<sub>2</sub>)<sub>2</sub>-[1,2,3-triazole]-CH<sub>2</sub>-**Big** | - 0.67 | 0.68 ± 0.06 |

<sup>a</sup> Big = Biguanide ; <sup>b</sup> cLogP were predicted by ChemDraw Professional Software (molecules were represented in their monoprotonated state to mimic physiological conditions) ; <sup>c</sup> The data are plotted as mean ± SD of 3 independent replicates ; <sup>d</sup> Met. = Metformin
See section “Material and Methods” in Supporting Information

Metformin is the most hydrophilic compound, with a cLogP of −1.63. However, this hydrophilicity poses a limitation, hindering its diffusion across biological membranes. Metformin’s transport relies on organic cationic transporters, ultimately impacting its activity (EC_50_ ∼ 0.85 mM). In contrast, E3 ligands **8** and **27**, with cLogP values of 0.40 and 2.10, respectively, depend predominantly on their permeability and diffusion across membranes. This heightened membrane permeability likely contributes to their higher antiproliferative activity, with E3 ligand **8** exhibiting an EC_50_ of approximately 0.17 mM, and E3 ligand **27** displaying an EC_50_ of around 0.34 mM. The varying hydrophobicity and permeability characteristics underscore the intricate balance between these factors and their consequential impact on the observed biological activities. In most instances, the conjugation of E3 ligands to the biguanide motif resulted in a higher partition coefficient compared to metformin, except for PROTAC **20**, which, due to its polar chain, exhibited increased hydrophilicity.Similarly PROTAC **16** demonstrated a cLogP value quite similar to that of metformin. Consequently, PROTACs **20** and **16**, being either more hydrophilic or comparably membrane-permeable to metformin, exhibited similar or restricted membrane permeability, reflected by EC_50_ values greater than or equal to that of metformin. Correspondingly, compounds with cLogP values less than 0 displayed the highest EC_50_ values, surpassing 0.34 mM. Conversely, the most apolar PROTACs, namely **11, 12**, and **29**, exhibited the highest activity against KP4, aligning with the trend that increased permeability can enhance compound efficacy. Although the study highlighted a relationship between cytotoxic activity and permeability, the median inhibitory concentration ranges of the synthesized PROTAC-Biguanides were not significantly distinct from those of their controls, namely metformin and the ligands CRBN_L_ and VHL_L_. Given these observations, it becomes imperative to delve into the mechanism of action of these compounds to discern whether their activity stems from the biguanide moiety, the E3 enzyme ligand, or if additional mechanisms, such as ternary complex formation, are at play.

## Mechanism of Action

The conjugation of the biguanide moiety with an E3 ligand introduces a duality in function, where the compound can either operate as a PROTAC, inducing the formation of a ternary complex leading to target protein degradation, or operate as one of its ligands, binding solely to a single target. To ascertain whether the synthesized compounds could effectively engage with the biguanide target, their capacity to activate the AMPK pathway was investigated. This pathway is known to be activated by phosphorylation in response to the administration of biguanidium salts. In KP4 cells after a 24-hour treatment with all Biguanide-PROTACs and their controls, it was observed that compounds **11** and **12**, could significatively induce AMPK phosphorylation at the concentrations studied (Fig 3A).

**Fig 3.**
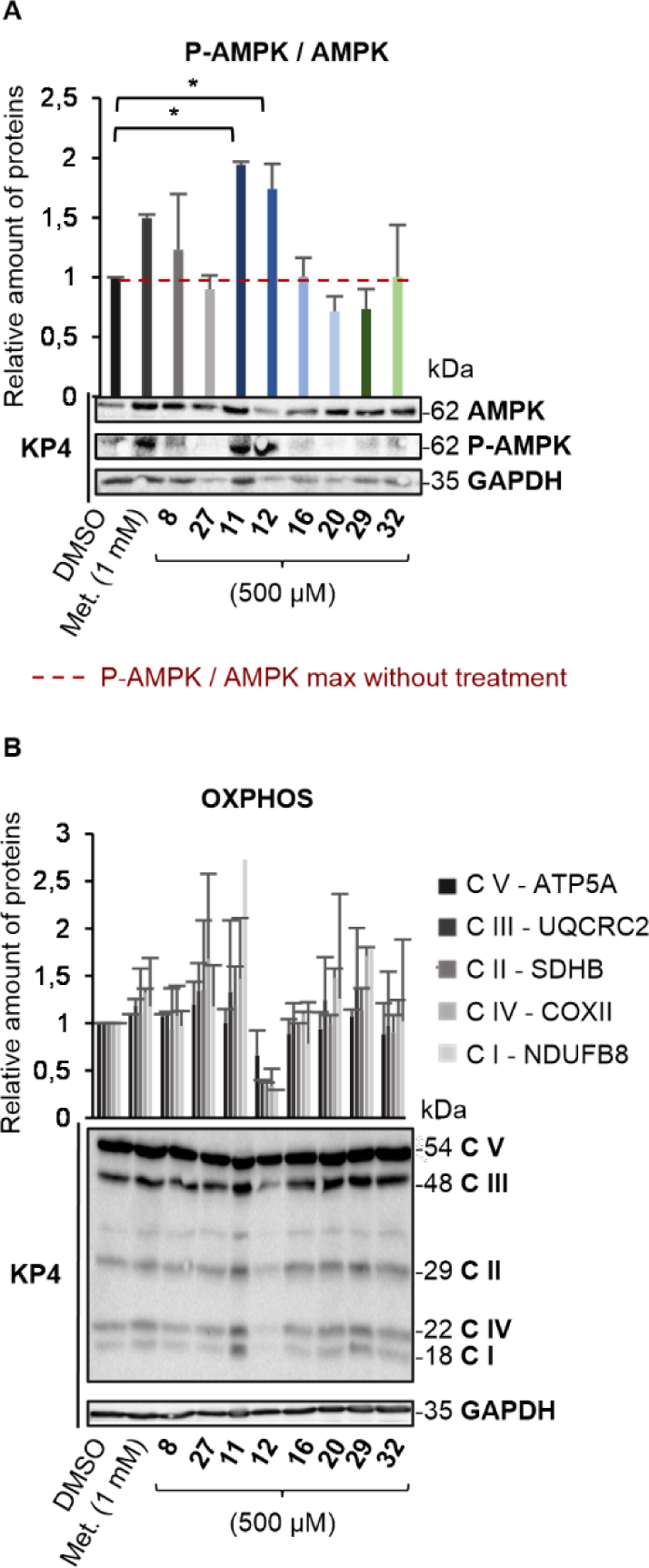
Biguanide-PROTACs effect on P-AMPK and OXPHOS proteins levels. (A) Immunoblot and relative quantification of P-AMPK and AMPK levels. (B) Immunoblot and relative quantification of OXPHOS proteins levels; both in KP4 after 24 h of treatment. ^*^p ≤ 0.05 (ANOVA).

This suggests that these particular compounds operate via their biguanide function, triggering AMPK activation. On the other hand, the activity of the remaining compounds may stem from their exclusive interaction with the E3 enzyme, reinforcing the hypothesis that their mode of action differs from that of metformin and compounds **11** and **12**.

To delve deeper into the mode of action, the levels of OXPHOS proteins from complexes I to V were measured by immunoblot. Notably, the relative quantities of these proteins did not exhibit a significant decrease after treatment with the various compounds, with the exception of compound **12** (Fig 3B). This observation hints at the complexity of the compounds’ interactions within the cellular machinery, prompting further exploration into the specific molecular pathways influenced by each compound.

The experiment was subsequently repeated with varying concentrations of these compounds, revealing a subtle yet noteworthy disturbance in proteins COXII (Complex IV) and NDUFB8 (Complex I) at concentrations of 250 µM and above (Fig 4). This observed alteration in protein levels could potentially be attributed to the degradation induced by Biguanide-PROTAC **12**. In this scenario, the proximity between the ubiquitin ligase E3 and mitochondrial proteins may have facilitated the ubiquitination of the target proteins. Given that compound **12** was the sole compound exhibiting activity, it is plausible to hypothesize that its hydrophobic linker might contribute to its accumulation in the mitochondria. This accumulation could then lead to a degradation of proximal proteins, providing a potential explanation for the observed alteration in the levels of COXII and NDUFB8.

**Fig 4.**
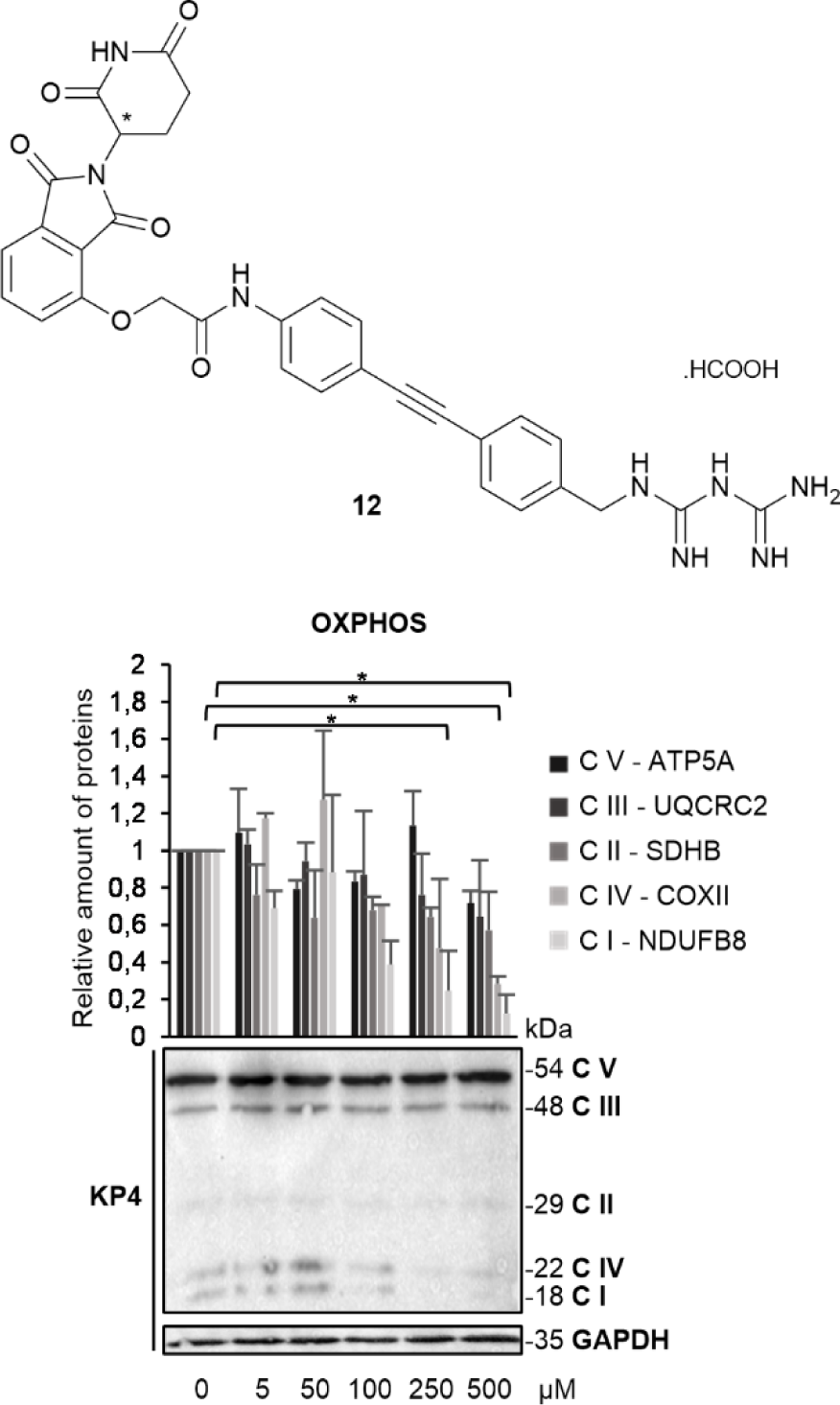
Effect of compound 12 on OXPHOS proteins levels. Immunoblot of OXPHOS proteins in KP4 after 24 h of treatment and relative quantification of protein levels. ^*^p ≤ 0.05 (ANOVA).

Nevertheless, it is difficult to identify with certainty a specific target for biguanides. Indeed, several studies have shown that mitochondrial respiratory chain complexes can be arranged in different ways: in isolated form or as supercomplexes.(42) In the latter case, the interaction of PROTAC **12** with one of these supercomplexes could lead to protein degradation of several assembled complexes. This insight raises intriguing questions about the specificity and selectivity of these compounds in engaging with cellular targets, underscoring the need for a comprehensive understanding of their mechanism of action for further therapeutic development. Moreover, the motif incorporated into compound **12**, known as phenylethynylbenzylebiguanide or PEB-Biguanide (Ph-C≡C-Bn-Biguanide), has been characterized as an amphiphilic cation with the ability to diffuse through phospholipidic membranes, as previously described.(15) The heightened antiproliferative properties of this motif, when compared to metformin, were attributed to its facilitated insertion into mitochondria. Mechanistic insights gleaned from studies on compound **12** suggest that the incorporation of this structure into more complex molecules give them the same properties. Thus it appears that our compound’s increased efficacy compared to metformin’s may stem not only from its enhanced cellular transport (see cLogP, Table 1) but also from its specific accumulation within mitochondria that enables proteins levels perturbation.

## Conclusion

The conjugation of the biguanide motif with various ligands of the E3 enzyme, incorporating distinct spacers between the two motifs, yielded a series of synthesized compounds. The impact of these compounds on the proliferation and viability of pancreatic cancer cells (KP4) facilitated the establishment of a correlation between the permeability of the active ingredient and its median inhibitory concentration. This foundational work laid the groundwork for exploring a structure-activity relationship, providing valuable insights into the factors influencing the compounds’ efficacy.

Further investigations into the mechanism of action of these Biguanide-PROTACs revealed that compounds **11** and **12** behaved akin to biguanides concerning the activation of AMPK. However, only compound **12** demonstrated the ability to alter levels of mitochondrial proteins, notably complexes I and IV. This distinctive observation suggests that the hydrophobic structure of compound **12** might facilitate its accumulation within mitochondria, thereby enhancing its antiproliferative properties. Looking ahead, the development and expansion of the library of Biguanide-PROTACs present an enticing prospect. This extension has the potential to yield new insights into the mechanism of action of biguanides, contributing to a deeper understanding of their targets, while concurrently enabling the creation of more potent anticancer agents. The ongoing exploration of this compound library promises to unveil novel therapeutic avenues, making significant strides in the quest for effective cancer treatments.

## Author Contributions

A.R.S and G.F. conceived the project, wrote funding grants and received funding for the project; J.V. performed the chemical and biological experiments; V.G. performed and supervised the biological experiments; J.V. wrote and V.B., G.F. and A.R.S edited the manuscript.

---

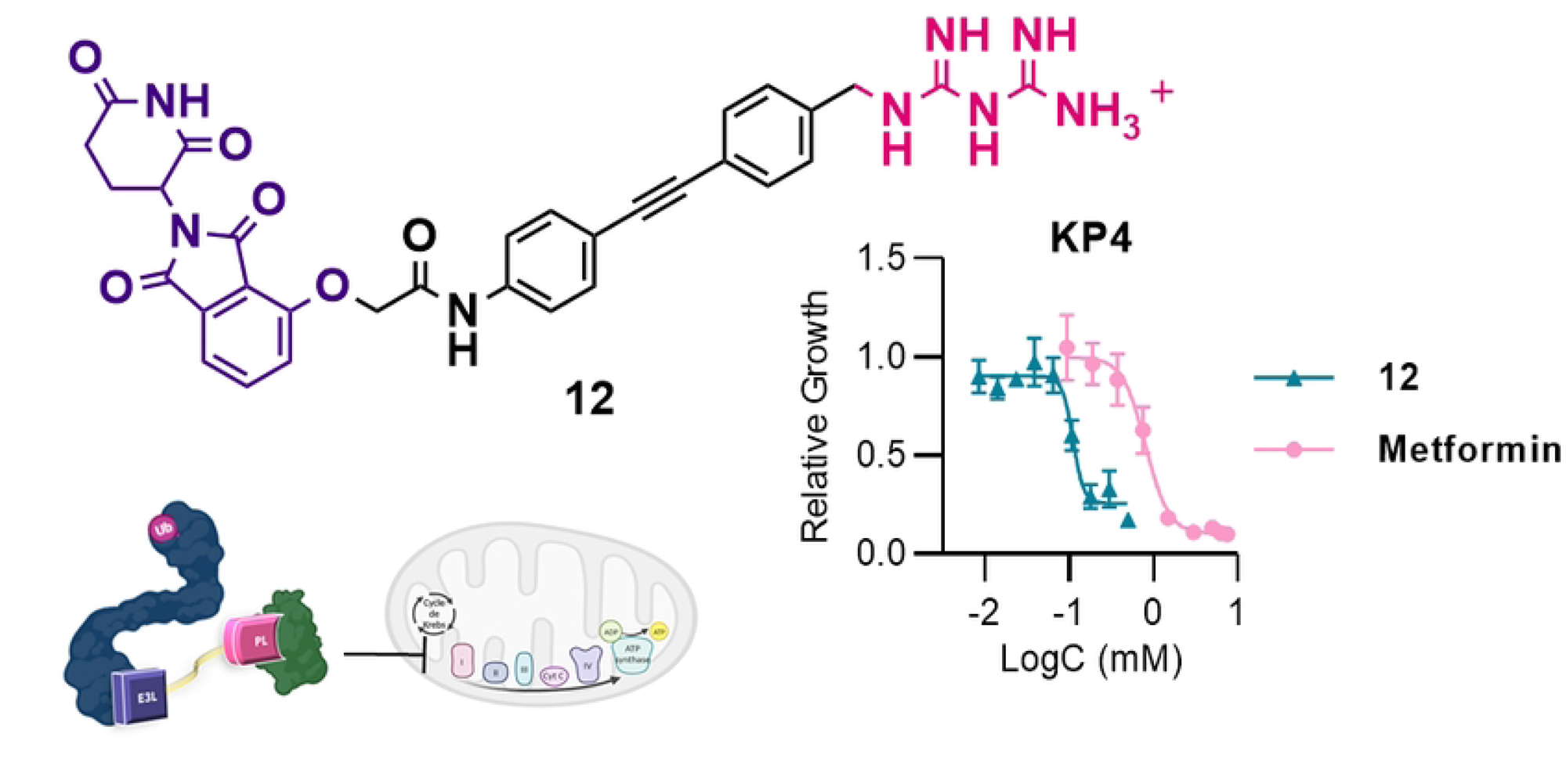

